# Self-fertilization and 1 inbreeding limit the scope for sexual antagonism

**DOI:** 10.1101/009365

**Authors:** Samuel J. Tazzyman, Jessica K. Abbott

## Abstract

Sexual antagonism occurs when there is a positive intersexual genetic correlation in trait expression but opposite fitness effects of the trait(s) in males and females. As such, it constrains the evolution of sexual dimorphism and may therefore have implications for adaptive evolution. There is currently considerable evidence for the existence of sexually antagonistic genetic variation in laboratory and natural populations, but how sexual antagonism interacts with other evolutionary phenomena is still poorly understood in many cases. Here we explore how self-fertilization and inbreeding affect the maintenance of polymorphism for sexually antagonistic loci. We expected *a priori* that selfing should reduce the region of polymorphism, since inbreeding reduces the frequency of heterozygotes and speeds fixation. Although this expectation was supported, our results show that there is an interactive effect between the degree of selfing and dominance such that those segregating sexually antagonistic loci that do exist are more likely to be partially dominant. In addition, inbreeding effects may influence population persistence and genomic location of sexually antagonistic loci in separate-sexed organisms.

## Introduction

Sexual antagonism occurs when there is a positive intersexual genetic correlation in trait expression but opposite fitness effects of the trait(s) in males and females (Bonduriansky & Chenoweth, 2009; it is also known as intralocus sexual conflict when the intersexual genetic correlation is for the same trait in both sexes). As such, it constrains the evolution of sexual dimorphism and may therefore have implications for adaptive evolution. Sexual antagonism has often been considered a relatively transient phenomenon which will eventually be resolved by the evolution of sex-specific modifiers (Stewart *et al*., 2010), but recent research suggests that pleiotropic effects can constrain the evolution of sex-limitation (Mank & Ellegren, 2009), and that even the evolution of sexual dimorphism need not resolve the conflict (Cox & Calsbeek, 2009; Harano *et al*., 2010). Some authors have even suggested that sexual antagonism is inevitable in any species with separate sexes (Connallon & Clark, 2014). The study of sexual antagonism has now matured to a point where its taxonomic ubiquity and potential importance in natural populations can no longer be questioned (Bonduriansky & Chenoweth, 2009; Cox & Calsbeek, 2009). The time has therefore come when we can no longer study sexual antagonism as an isolated phenomenon; it must now be integrated with broader evolutionary theory. To this end, we have constructed a model that investigates how self-fertilization and inbreeding affect the maintenance of polymorphism at sexually antagonistic loci.

Our motivation was to begin formalizing recent speculations about the nature and relevance of sexually antagonistic variation in hermaphroditic species (Abbott, 2011; Bedhomme *et al*., 2009). All else being equal, sexually antagonistic alleles in hermaphrodites should exhibit the same dynamics as autosomal sexually antagonistic loci in separate-sexed organisms. This implies that selection must be strong and approximately equal in magnitude across the sexes in order for polymorphism to be maintained (Kidwell *et al*., 1977). However hermaphrodites have a number of properties that set them apart from species with separate sexes, one of which is the fact that many hermaphrodites are partially or completely selfing (Goodwillie *et al*., 2005; Jarne & Auld, 2006). Selfing should reduce the region of polymorphism, since inbreeding reduces the frequency of heterozygotes and speeds fixation (Lynch & Walsh, 1998). One might therefore expect *a priori* that partially selfing hermaphrodites should exhibit lower levels of sexually antagonistic genetic variation compared to separate-sexed species or obligate outcrossers. However recent work by Fry (2009) demonstrates that dominance effects can have considerable influence on the region of parameter space permitting polymorphism. We therefore extended the framework developed by Kidwell *et al*. (1977) and Fry (2009) to investigate how proportion selfing and dominance interact to affect the maintenance of sexually antagonistic loci. We found that although inbreeding reduces the region of parameter space permitting polymorphism overall, it can offset some of the effects of dominance demonstrated by Fry (2009).

## Model

Our model is based on a classic framework for the investigation of sexually antagonistic alleles (Kidwell *et al*., 1977). Our population is made up of a large number of diploid hermaphroditic individuals. We focus on a single locus, at which there are two alleles, denoted *A* and *a.* This means that every individual is one of three genotypes: *AA, Aa,* or *aa.* Each genotype confers a different fitness; there are assumed to be no other differences between individuals.

We model generations as being discrete and non-overlapping. Within each generation, the life cycle goes as follows. We first census the genotypes in the population, and denote the frequency of genotype *AA* by p, and the frequency of genotype *aa* by q. Then the frequency of heterozygote *Aa* types is 1 *-p - q.*

After censusing, random mating occurs. A proportion *F* of matings are self-fertilisation, while the remaining 1 - *F* matings are outbreeding events. For the self-fertilisation events there is no effect of genotype on offspring production. The genotypes of the offspring from self-fertilisation will depend on the parental genotype. Homozygous *AA* or *aa* individuals will produce their own genotypes for offspring, while heterozygous *Aa* individuals will have 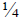 of their offspring of genotype *AA,* 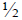 genotype *Aa*, and 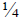 genotype *aa* (Table 1). Thus the frequency of each genotype in the next generation due to offspring from inbreeding events is as follows. 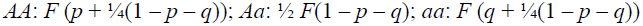.

**Table 1.** Offspring genotype distributions when inbreeding

| Parental genotype | Offspring genotype |
| --- | --- |
| $AA$ | $AA$ |
| $Aa$ | $\frac{1}{4} AA, \frac{1}{2} Aa, \frac{1}{4} aa$ |
| $aa$ | $aa$ |

Because we are assuming random mating we can model each outbreeding event as being the result of the combination of sperm and eggs from randomly drawn individuals, with the probability of drawing a given genotype in a given sex role being proportional to the frequency of that genotype, and to its fitness in that sex role. Fitness differs across genotypes and across each sex role, as summarised in Table 2. The *A* allele is female-beneficial, male-deleterious, so that bearing an *A* allele makes a hermaphroditic individual better at the female role but worse at the male role. Conversely, the *a* allele is female-deleterious, male-beneficial, so that bearing an *a* allele makes an individual worse at the female role but better at the male role. These deleterious and beneficial effects are summarised by the parameters *sf* and *s_m_,* which represent the selection coefficients, and *hf* and *h_m_,* which represent the dominance coefficients (Table 2).

**Table 2.**
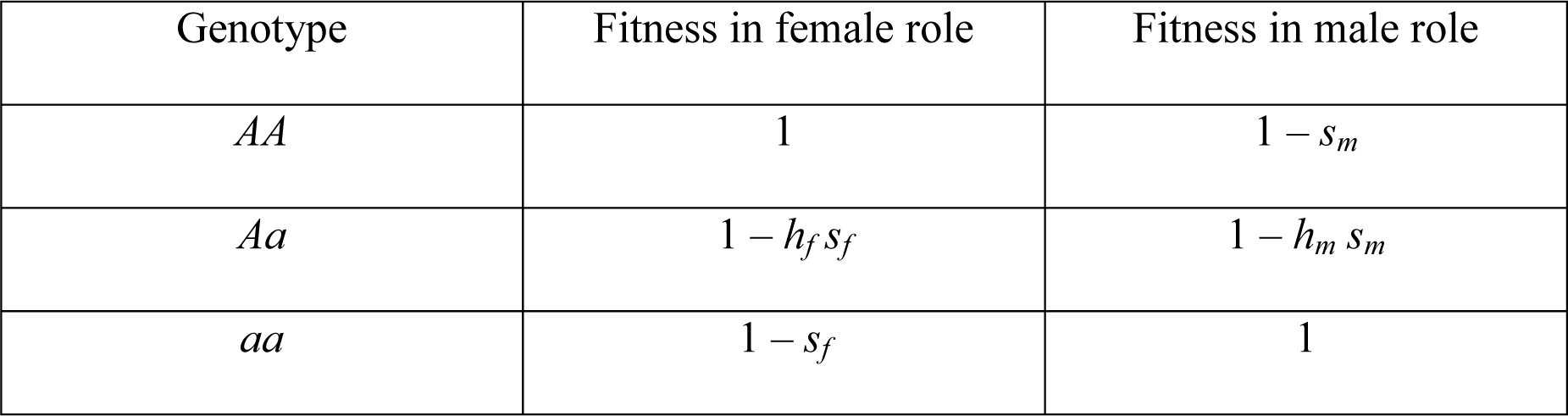
Fitness in different sex roles when outbreeding

| Genotype | Fitness in female role | Fitness in male role |
| --- | --- | --- |
| $AA$ | 1 | $1 - s_m$ |
| $Aa$ | $1 - h_f s_f$ | $1 - h_m s_m$ |
| $aa$ | $1 - s_f$ | 1 |

Since we assume that there are a large number of outbreeding events, the frequency of each genotype in the next generation due to offspring from inbreeding events is as follows. The frequency of *AA* individuals is

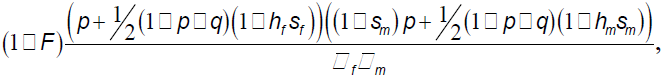

the frequency of *aa* individuals is

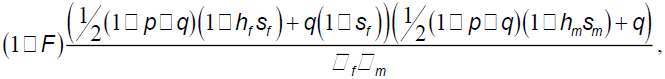

where 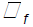 and 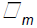 are respectively the mean fitness in the female and male roles,

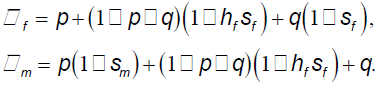

The frequency of *Aa* individuals is the balancing expression so that these three frequencies add to (1 - *F).*

Putting together both self-fertilisation and outbreeding events, we can derive expressions for the change in frequency of genotypes *AA* and *aa* from one generation to the next, denoted *Δp* and *Δq* respectively, as

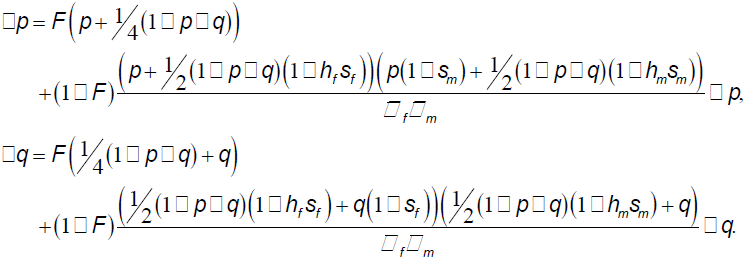

Using equations (1) we can establish whether the *A* and *a* alleles are protected from extinction when rare, and consequently whether polymorphism is protected or not (Appendix), depending on the values of parameters F, *sf, s_m_, hf,* and *h_m_.* When *F* = 0, we recover the classic results for this model (Fry, 2009; Kidwell *et al*., 1977). Therefore in our analysis here we focus on the effect of *F* on the region admitting polymorphism.

## Results

We can use (1) to calculate expressions *a* and *A* corresponding to protection when rare of *a* and *A*, respectively. The female-deleterious, male-beneficial allele *a* is protected from extinction where rare if

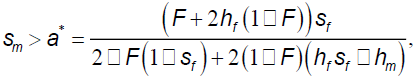

while the female-beneficial, male-deleterious allele *A* is protected from extinction when rare if

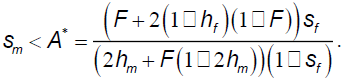

We want to know the effect of *F* on these thresholds. Thus, we calculate

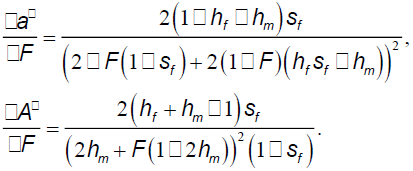

If δ*a*\*/δ*F* < 0, and δ*A*\*/δ*F* > 0, then increasing *F* increases the region of parameter space that leads to polymorphism. This is true if *hf* + *h_m_* > 1, and false if *hf* + *h_m_* < 1. If *hf* + *h_m_* = 1 (for example if both alleles are additive in their effects), changes in the proportion of inbreeding make no difference to whether or not there is a polymorphism.

Interestingly, this result relates to a previous finding, which showed that in the absence of inbreeding, the higher the value of *hf* + *h_m_,* the smaller the region of parameter space admitting polymorphism (Fry, 2009). For any fixed value of *F* this remains the case in our model. However, this dominance effect is weakened by inbreeding, because inbreeding results in fewer heterozygotes, and consequently the effect of *hf* and *h_m_* is weakened (Figure 1). The results when *F* = 1 are identical to the case when *hf* = *h_m_* = 0.5.

**Figure 1.**
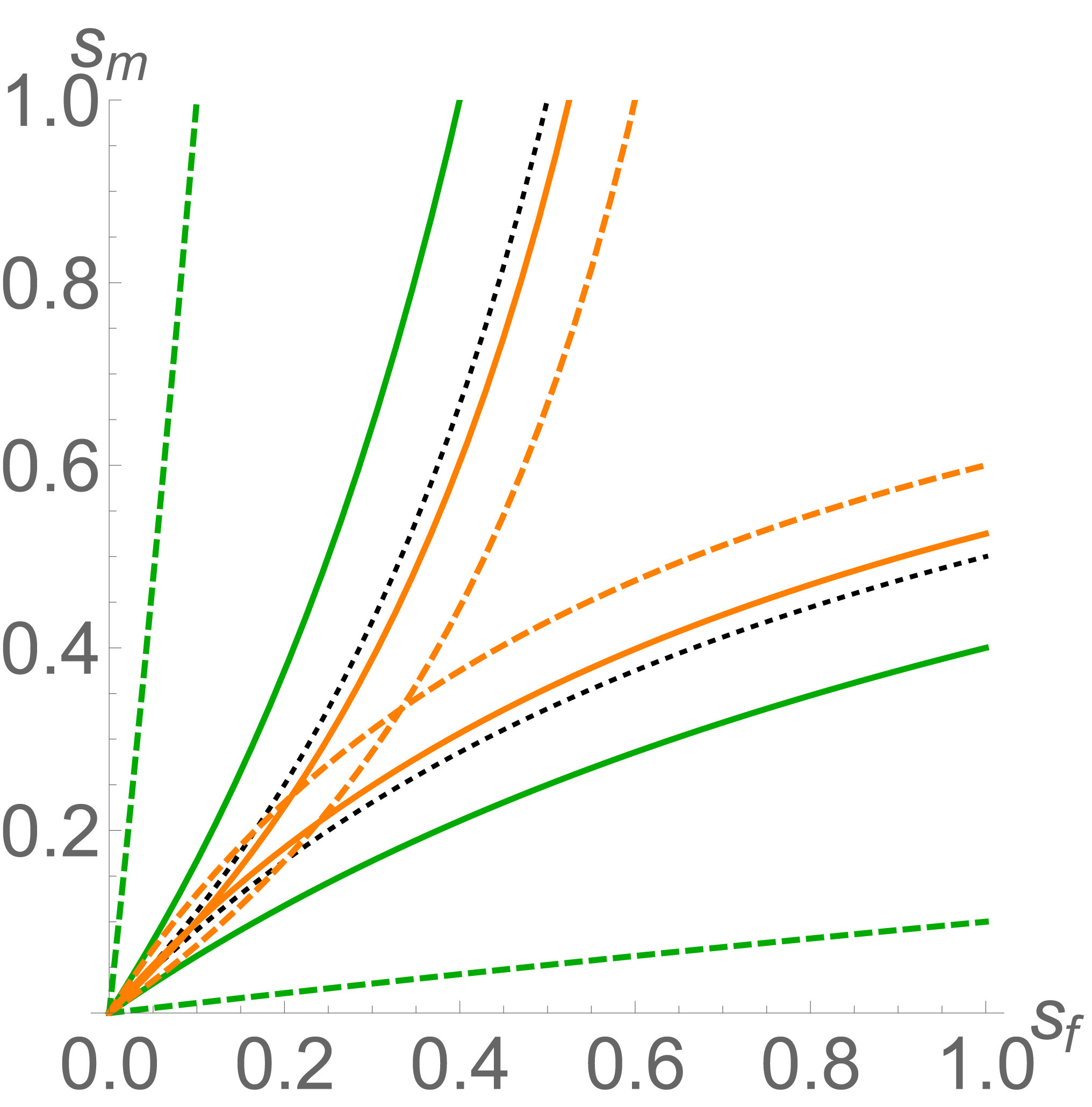
The effect of inbreeding and dominance on the maintenance of sexually antagonistic polymorphisms in hermaphrodites. The area between two matching curves is where polymorphism is admitted. The green curves correspond to the case where *hf* = *hm* = 0.1, so that the allele that is deleterious in each sex is partially recessive in that sex. The orange curves correspond to the case where *hf* = *h_m_* = 0.6, so that the allele that is deleterious in each sex is partially dominant in each sex. The dashed curves represent where *F* = 0, the situation where there is no inbreeding. The solid curves represent the case where *F* = 0.75, so that three quarters of matings are self-fertilisation. For the green curves, this results in a smaller area of polymorphism, while for the orange curves, it results in a larger area of polymorphism. The black dotted line is the asymptotic limit *F* = 1. It exactly corresponds to the case in which *F* = 0 and *hf* = *h_m_* = 0.5.

For weak selection (e.g. parameter values 0 < *sf, s_m_* < 0.1) there is very little scope for polymorphism when *hf* + *h_m_* > 1 (Figure 2, see also Fry (2009)) regardless of the value of *F.* However, when *hf* + *h_m_* < 1, the range of parameter values for which there can be a sexually antagonistic polymorphism due to weakly selected alleles is severely curtailed by self-fertilisation (Figure 2).

**Figure 2.**
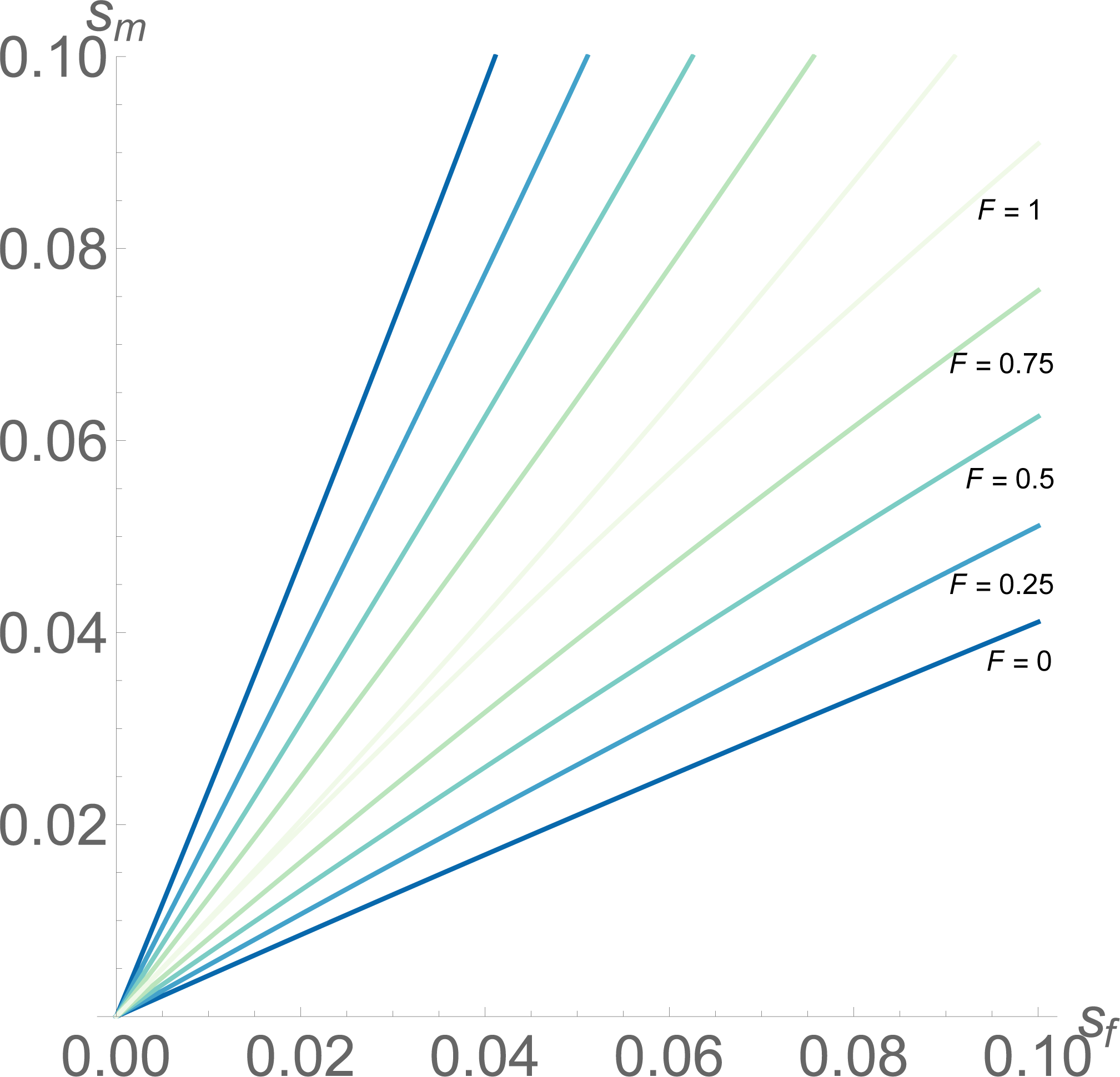
The effect of inbreeding on the maintenance of weakly sexually antagonistic polymorphisms. The region between any two matching-coloured lines admits a stable sexually antagonistic polymorphism. The pairs of matching lines correspond to the cases where ***F*** = 0, 0.25, 0.5, 0.75, and 1, respectively, as marked. Here ***hf*** = ***h_m_*** = 0.3; as inbreeding increases, the region admitting polymorphism decreases in size, to the limiting case where ***F*** = 1. For values of ***hf*** + ***h_m_*** > 1, the region admitting polymorphism is contained within the region for ***F*** = 1, and consequently for these dominance parameters there is very little scope for polymorphism under weak selection.

## Discussion

In hermaphrodites, the ability to self-fertilise will affect the maintenance or otherwise of sexually antagonistic polymorphisms. We expected *a priori* that selfing should reduce the region of polymorphism, since inbreeding reduces the frequency of heterozygotes and speeds fixation (Lynch & Walsh, 1998). Although this expectation was supported, our results show that there is an interactive effect between the degree of selfing and dominance. In species in which there is no self-fertilisation (e.g. separate-sexed organisms and obligate outcrossers) the more an allele that is deleterious to one sex is dominant in that sex, the smaller the region of parameter space that will admit polymorphism (Fry, 2009). However, this effect is weakened by self-fertilisation (Figure 1), so that in partially selfing hermaphrodites we would expect more dominant sexually antagonistic alleles remaining at polymorphism (and fewer recessive alleles) than if there were no selfing. In particular, for weakly selected sexually antagonistic alleles which are on average partially recessive in their deleterious state, the range of parameter space allowing for a polymorphic equilibrium is strongly restricted in the case where there is inbreeding (Figure 2); if the alleles are on average partially dominant in their deleterious state, the region of parameter space allowing for polymorphism is increased by inbreeding, but remains small. Overall, therefore, the more self-fertilisation occurs in hermaphrodites, the fewer sexually antagonistic polymorphisms we would expect (assuming that most sexually antagonistic selection is weak).

Our result is clear and simple, and shows how the ability to self-fertilise can have a strong effect on the genetics of a species. However, the effect of inbreeding is not strong enough to completely cancel out the effect of dominance: for a given frequency of inbreeding, it will still be the case that the more dominant the alleles are in their deleterious context, the smaller the region of parameter space in which they can exist at polymorphism.

It is of course well-established that selfing can lead to inbreeding depression, but that in habitually selfing organisms the benefits of selfing should outweigh the costs of inbreeding depression (Goodwillie *et al*., 2005). We have therefore not explicitly modelled inbreeding depression, and have assumed that selfed gametes do not experience selective effects of the sexually antagonistic alleles. The logic behind this assumption was that because selfing is usually considered a form of reproductive assurance (Goodwillie *et al*., 2005), this assurance will be ineffective if selfed gametes are subject to the same selection pressures as outcrossed gametes. This assumption is realistic if most of the cost of outcrossing is due to extrinsic factors, such as sexual conflicts with the mating partner (Anthes & Michiels, 2007; Koene, 2006; Koene *et al*., 2005) or energetic or predation costs of finding a mate (Jennions & Petrie, 1997). It becomes less realistic if the sexually antagonistic alleles cause intrinsic fitness differences (e.g. poor survival of gametes). Sperm (or pollen) limitation is unlikely to be a major limiting factor in fecundity when selfing (but see Hodgkin & Barnes, 1991), but it is not unlikely that mutations affecting egg quality/survival would have an effect even on the production of selfed offspring. If sperm are accompanied by toxic seminal fluid used in sperm competition when outcrossing, then this could also contribute to lower egg survival, even when selfing (Koene *et al*., 2010; Schärer *et al*., 2014). On the other hand, positive selection of alleles with a negative impact on egg production should be weak in species with high levels of selfing, because there is limited scope for fitness gain via sperm donation during outcrossed matings. We therefore believe that this assumption is reasonable.

It is also worth noting that our model, although originally constructed with hermaphrodites in mind, is equally applicable to separate-sexed organisms with respect to inbreeding instead of selfing (Appendix). This generates some interesting predictions, especially for populations with high levels of inbreeding, such as island populations (e.g. Grant *et al*., 2003), or populations with low dispersal levels due to habitat fragmentation (e.g. Andersen *et al*., 2004).

Firstly, species or populations with high levels of inbreeding should exhibit reduced levels of sexual antagonism compared to outbred populations, because the gain in polymorphism in dominant alleles is more than offset by the loss in polymorphism in recessive alleles, for a given level of inbreeding (Figure 1). A recent model suggests that sexual antagonism and demography can interact to cause extinction of populations located in patches that are beneficial to male fitness and detrimental to female fitness (Harts *et al*., 2014). This is because populations collapse if there are too few reproducing females. However populations which are declining in numbers should also become more inbred, as a result of the decreasing effective population size. Our results therefore suggest that the negative effect of sexually antagonistic alleles on population growth may be mitigated by inbreeding. This could reduce the chance of population collapse.

Secondly, we should expect a higher proportion of sexually antagonistic alleles to be located on the X-or Z-chromosomes in inbred populations relative to outbred populations. Heteromorphic sex chromosomes have long been held to be likely hotspots for sexual antagonism. In a seminal paper, Rice (1984) argued that the X-chromosome should harbour increased levels of sexually antagonistic genetic variation because male-benefit loci that are recessive in females will be expressed in hemizygous males, but largely escape counter-selection in females at low to intermediate frequencies. Conversely, dominant female-benefit loci will also be more common on the X than on the autosomes, despite their deleterious effect in males, because the X spends more time in females than the autosomes (2/3 versus ^) and therefore experiences stronger total female-specific selection. Our results suggest that the heteromorphic sex chromosomes should harbour higher levels of sexually antagonistic genetic variation in inbred populations, independent of the direction of the fitness effect (i.e. female-benefit/male-detriment versus male-benefit/female-detriment). This is because any X-or Z-linked locus that is not completely recessive in the homogametic sex will be partially dominant overall (i.e. *hf* + *h_m_* > 1 will always hold true when *h*_*homogametic*_ > 0 because *h*_*heterogametic*_ = 1), and therefore subject to an increased range of polymorphism with increasing inbreeding level.

In sum, we show that although inbreeding reduces the region of parameter space permitting polymorphism overall, it can offset some of the effects of dominance demonstrated by Fry (2009). This means that although hermaphrodites with high levels of inbreeding are unlikely to harbour significant sexually antagonistic genetic variation, those segregating sexually antagonistic loci that do exist are more likely to be partially dominant. In addition, inbreeding effects may influence population persistence and genomic location of sexually antagonistic loci in separate-sexed organisms.

## Acknowledgements

SJT was supported by funding from the European Research Council under the 7th Framework Programme of the European Commission (PBDR: Grant Agreement Number 268540), and JKA was supported by funding from the Swedish Research Council.

## Appendix

### Stability of equilibria

Using equations (1) we can define the function *g[p,* q] = (Δ*p*, Δ*q*), defined for all possible values of*p* and *q* (i.e. on the standard 2-simplex). We know that g[1, 0] = (0, 0) (corresponding to fixation of the *A* allele), and g[0, 1] = (0, 0) (corresponding to fixation of the *a* allele). To determine whether either of these two equilibria are stable we consider the Jacobian matrix **J** of the function g,

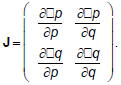

For each fixed point, we evaluate **J** and calculate its eigenvalues. If they are all negative for a given equilibrium point, that point is stable (thus if any of the eigenvalues are positive, the equilibrium point is unstable). If the equilibrium point at (1, 0) is unstable, then *a* is protected from extinction when rare (corresponding to the condition *s_m_* > *a\** given in the main text). If the equilibrium point at (0, 1) is unstable, then *A* is protected from extinction from rare (corresponding to the condition *s_m_* < *A\** given in the main text). If both alleles are protected from extinction when they are rare, then we have a protected polymorphism.

### Applicability of model to separate-sexed species

Although the model was constructed to consider hermaphrodites, it can also apply to separate-sexed species. Because separate-sexed species cannot self-fertilise, the definition of *F* as the proportion of self-fertilising events cannot be maintained. Instead, *F* is taken to be a measure of the additional probability with which an individual will mate with a partner sharing the same genotype at the *A*/*a* locus of interest (Appendix Table 1). Thus *F* can be seen as a measure of the level of inbreeding that is occurring in the population.

**Appendix Table 1:** Application of *F* to separate-sexed species

| Focal genotype | Mating partner's genotype | Probability of that partner genotype under random mating | Probability of that partner genotype with "self-fertilisation frequency" $F$ |
| --- | --- | --- | --- |
| $AA$ | $AA$ | $p$ | $p + F(1 - p)$ |
| | $Aa$ | $1 - p - q$ | $(1 - F)(1 - p - q)$ |
| | $aa$ | $q$ | $(1 - F)q$ |
| $Aa$ | $AA$ | $p$ | $(1 - F)p$ |
| | $Aa$ | $1 - p - q$ | $1 - p - q + F(p + q)$ |
| | $aa$ | $q$ | $(1 - F)q$ |
| $aa$ | $AA$ | $p$ | $(1 - F)p$ |
| | $Aa$ | $1 - p - q$ | $(1 - F)(1 - p - q)$ |
| | $aa$ | $q$ | $q + F(1 - q)$ |

